# Activity of prefrontal cortex serotonin 2A receptor expressing neurons is necessary for the head-twitch response of mice to psychedelic drug DOI in a sex-dependent manner

**DOI:** 10.1101/2024.05.21.595211

**Authors:** Annika B. Ozols, Jing Wei, Janet M. Campbell, Chengcheng Hu, Shenfeng Qiu, Amelia L. Gallitano

**Affiliations:** * Department of Basic Medical Sciences, University of Arizona College of Medicine – Phoenix, 425 N. 5^th^ St., Phoenix, AZ, 85004; Epidemiology and Biostatistics, University of Arizona Mel and Enid Zuckerman College of Public Health – Phoenix, 714 E Van Buren St #119, Phoenix, AZ, 85006

## Abstract

Serotonin 2A receptors (5-HT_2A_Rs) mediate the effects of psychedelic drugs. 5-HT_2A_R agonists, such as (-)-2,5-dimethoxy-4-iodoamphetamine hydrochloride (DOI), that produce a psychedelic experience in humans induce a head-twitch response (HTR) behavior in rodents. However, it is unknown whether the activity of 5-HT_2A_R expressing neurons is sufficient to produce the HTR in the absence of an agonist, or in which brain region 5-HT_2A_Rs control the HTR. Here, we use an optogenetic approach to examine whether activation of 5-HT_2A_R expressing neurons in the mouse prefrontal cortex (PFC) is sufficient to induce HTRs alone, or may augment the HTR produced by DOI, and if inhibition of these neurons prevents DOI-induced HTRs in mice. We crossed *Htr2a*-Cre mice to Cre-dependent optogenetic lines Ai32 (channelrhodopsin) and Ai39 (halorhodopsin) to selectively activate and inhibit (respectively) 5-HT_2A_R-expressing neurons in the PFC of adult mice. We found that optogenetic stimulation of PFC 5-HT_2A_R expressing neurons in the absence of an agonist does not increase HTRs in mice. In both male and female Ai32 mice that received vehicle, there was no difference in HTRs in mice that expressed *Htr2a*-Cre compared with control mice, indicating that optogenetic activation of 5-HT_2A_R+ cells in the PFC was not sufficient to produce HTRs in the absence of an agonist. In female mice, activation of PFC 5-HT_2A_R expressing neurons augmented the HTR produced by DOI. However, this result was not seen in *male* mice. In contrast, inhibition of 5-HT_2A_R expressing neurons in the PFC prevented the increase in HTR produced by DOI in male, but not in female, mice. Together, these findings suggest that activation of 5-HT_2A_Rs in the PFC is not sufficient to induce HTRs in the absence of a 5-HT_2A_R agonist but is necessary for induction of HTRs by a 5-HT_2A_R agonist in a sex-dependent manner.

## Introduction

Serotonin 2A receptors (5-HT_2A_Rs) mediate the effects of psychedelic drugs such as psilocybin, lysergic acid (LSD), and mescaline [1,2]. In addition, 5-HT_2A_Rs are implicated in the etiology and treatment of numerous psychiatric disorders, from anorexia and bulimia nervosa [3–5] to schizophrenia [6–14] and major depressive disorder [11,13,15–18]. There has been a recent resurgence of interest in the use of psychedelics as therapeutics to treat a range of mental health and substance use disorders [19–22]. Recent studies have shown that psilocybin, and in once case LSD [23], alleviates anxiety related to a life-threatening cancer diagnosis [24–26] and induces a rapid, and long lasting, remission of symptoms in patients with refractory depression [27–29]. However, the mechanisms underlying these responses remain unknown.

5-HT_2A_R agonists, such as (-)-2,5-dimethoxy-4-iodoamphetamine hydrochloride (DOI), induce a distinctive behavior in rodents called the head-twitch response (HTR) [30,31]. The HTR has been extensively used as a behavioral assay to assess the physiological function of 5-HT_2A_Rs [32–37]. This behavioral response is dependent on 5-HT_2A_Rs as evidenced by studies showing that 5-HT_2A_R antagonists prevent induction of HTR [30,38,39] and the fact that HTR is absent in *Htr2a*-deficient mice [35,40]. Moreover, the HTR distinguishes 5-HT_2A_R agonists that produce a psychedelic experience in humans from those that do not have this effect [35]. This is important since the psychedelic experience may be an essential element in the therapeutic effect of drugs like psilocin, the active metabolite of psilocybin [41]. However, the brain region in which 5-HT_2A_Rs mediate HTRs in rodents remains unclear.

Human studies using 5-HT_2A_R ligand Positron Emission Tomography (PET) tracers show binding to regions throughout the cortex [42–44]. Studies in rodents, including our prior RNAscope *in situ* hybridization results, as well as previous radioactive *in situ* hybridization and antibody staining studies, have shown that both *Htr2a* mRNA, which encodes the 5-HT_2A_R, as well as the receptor protein itself, display an anteroposterior gradient throughout the cortex [45–48]. While the specific region in which 5-HT_2A_Rs mediate HTR is not certain, regional injection of DOI into the PFC has been shown to induce HTRs in rats [38]. However, it is not known whether the activity of 5-HT_2A_R expressing neurons is sufficient to produce HTR in the absence of an agonist, or whether inhibition of these neurons in the PFC abrogates the effect of psychedelics. Here, we use an optogenetic approach to determine whether activation of 5-HT_2A_R-expressing neurons in the mouse prefrontal cortex (PFC) is sufficient to induce HTRs and if inhibition of these neurons prevents HTRs in mice following DOI administration.

## Materials and Methods

### Animals

*Htr2a*-Cre mice (B6.FVB(Cg)-Tg(Htr2a-cre)KM207Gsat/Mmucd) on a C57BL/6J background were obtained from the Mutant Mouse Resource and Research Center (MMRRC) [49,50]. *Htr2a*-Cre mice were crossed to the following lines: Ai14 Cre-dependent tdTomato (JAX stock #: 007914, [51]), Ai39 Cre-dependent halorhodopsin (JAX stock #: 014539, [52,53]), and Ai32 Cre-dependent channelrhodopsin (JAX stock #: 024109, [52]). Matched pairs of littermate *Htr2a*-Cre^+/-^;Ai39^+/-^and *Htr2a*-Cre^-/-^;Ai39^+/-^ as well as *Htr2a*-Cre^+/-^;Ai32^+/-^ and *Htr2a*-Cre^-/-^;Ai32^+/-^ were designated at weaning. Both male and female mice were used for all experiments. Animals were housed on a 12-hour light/dark cycle with *ad libitum* access to food and water. All study protocols were performed in compliance with the Institutional Animal Care and Use Committee (IACUC) of University of Arizona.

### Validate *Htr2a*-Cre expression

One to 2-month old *Htr2a*-Cre^+/-^;Ai14^+/-^ mice were sacrificed via isoflurane overdose, then perfused with 1x phosphate-buffered saline (PBS) followed by 4% paraformaldehyde (PFA). Brains were post-fixed in PFA for 24 hrs, transferred to 30% sucrose at 4°C until saturation, frozen and sectioned at 40µm on a sliding microtome (American Optical). Free-floating sections were washed briefly in 1x PBS then mounted with Vectashield containing DAPI (Vector Laboratories, Cat #: H-1200). Slides were imaged with fluorescent microscopy (Zeiss, Axio Imager M2) at 10x magnification in DsRed (150ms) and DAPI (150ms) channels. Qualitative analyses were performed via thorough review of tissue sections and in reference to anatomical regions described in The Allen Mouse Brain Atlas (Reference Atlas, Interactive P56, Coronal) [54,55].

### Whole-cell patch clamp

Whole cell patch clamp recording was conducted in prefrontal cortical neurons in acute brain slices. ∼1 month old *Htr2a*-Cre^+/-^;Ai39^+/-^ mice and *Htr2a*-Cre^+/-^;Ai32^+/-^ mice were sacrificed and brains were removed. 350-μm coronal PFC-containing slices were made on a vibratome (VT1200S; Leica) in ice-cold artificial cerebrospinal fluid (ACSF) (126mM NaCl, 2.5mM KCl, 26mM NaHCO3, 2mM CaCl2, 1mM MgCl2, 1.25mM NaH2PO4, and 10mM glucose saturated with 95% O_2_ and 5% CO_2_). Slices were incubated at 32°C for 30 min for recovery before being transferred to a perfusion chamber mounted on a microscope (BX-51WI; Olympus) equipped with a water immersion objective (60×, NA 0.9). Neurons were visualized by fluorescence of the EYFP reporter, expressed in the Ai39 and Ai32 lines under brightfield differential interference contrast and epifluorescence illumination. Neurons with a soma at least 50μm below the slice surface were selected for recording. Tight seals (2–10 GΩ) were obtained by applying negative pressure. The membrane was disrupted with additional suction, and the whole-cell configuration was obtained. Patch electrodes containing internal solution: 130mM K-gluconate, 10mM HEPES, 4mM KCl, 0.3mM GTP-Na, 4mM ATP-Mg, 2mM NaCl,1mM EGTA and 14mM phosphocreatine (pHLJ7.2, 295–300LJmOsm) were used for action potentials recording. The resistance of patch electrode was ∼LJ4.0 MΩ. A small depolarizing current was applied to adjust the inter-spike potential to ∼ 60 mV. A MultiClamp 700B patch clamp amplifier (Molecular Devices), was used to condition and amplify the signal. Recorded signals were low-pass filtered at 1 kHz (voltage clamp) or 10kHz (current clamp) and digitized at 20 kHz using a Digidata 1440A board under control of pClamp 10.6 (Molecular Devices).

### Fiber optic cannulae implantation

Stereotaxic surgery was performed in 3.5-5 months old mice under 1-3% isoflurane anesthesia (VetOne). A circular incision was made to remove a portion of the superior scalp. The skull surface was cleaned via micro curette and dried with a sterile cotton tip. Fiber optic cannulae (RWD Life Science, Cat#: R-FOC-L200C-22NA, ID 1.25mm Ceramic Ferrule, 200um Core, 0.22NA, length 2.0mm) were bilaterally implanted into the PFC using the following coordinates at the final location relative to bregma: AP +2.1mm, ML ± 0.48mm, and DV −1.6mm, with lateral to medial angle of 10°. The fiber optic cannulae were secured with Loctite Ultra Gel Control super glue and C&B Metabond dental cement (Parkell). A steel wire loop, approximately the gauge of a paper clip, was mounted to the posterior surface of the skull with dental cement for removable attachment of a neodymium magnet (6.35 mm x 6.35 mm x 3.175 mm). All animals were allowed a 2-week recovery period prior to testing. Cannula placements were validated to be within ± 0.3mm of the AP target in all animals by post-mortem histology.

### Drug preparation and administration

(-)-1-(2,5-dimethoxy 4-iodophenyl)-2-aminopropane hydrochloride (DOI, Sigma-Aldrich) was and dissolved in sterile water at a concentration of 1mg/mL; serially diluted to 0.3mg/mL and 0.1mg/mL in sterile water, and aliquots of all three concentrations as well as sterile water (vehicle) were stored at −20°C until the day of testing. Drug or vehicle was administered via intraperitoneal (i.p.) injection at a final concentration of 0.0mg/kg, 0.1mg/kg, 0.3mg/kg, or 1.0mg/kg of body weight.

### Detection of head-twitch response

Behavioral testing was performed on 4-8.5 month old mice habituated to handling for 5min/day x 3 days. 24 hours prior to HTR experiments, mice were habituated to optogenetic patch cords for 10min. HTRs of Ai39^+^ mice were tested following 0.0mg/kg and 1.0mg/kg doses of DOI. Ai32^+^ mice underwent HTR experiments at 0.0mg/kg, 0.1mg/kg, 0.3mg/kg, and 1.0mg/kg doses of DOI. Animals were tested with each dose in randomized order with a washout period ≥ 3 days between tests. The half-life of DOI in mouse whole blood and forebrain is 1.90 ± 0.07 hrs and 1.52 ± 0.04 hrs, respectively [40]. Immediately following administration of either vehicle or DOI (i.p.), mice were placed in an 11cm diameter glass beaker surrounded by a copper wire coil (∼500 loops, 30 AWG) for 30 min; only the last 15 min of each recording session were analyzed. Both terminals of the copper coil were connected to a phono preamplifier (Pyle PP444), which was connected to a myDAQ (National Instruments) data acquisition system. The myDAQ was controlled via MATLAB software (MathWorks, version R2022a, with Data Acquisition Toolbox Support Package) which collected the amplified voltage signal at 1000Hz. To mitigate unwanted disturbances in electrical signal, the magnetometer was placed in a copper Faraday cage.

Automated head-twitch detection relied on a MATLAB script that analyzed individual events over the course of the recording session. The MATLAB script was based upon a previously validated automated head-twitch detection system by de la Fuente Revenga and colleagues [40]. The script rejected frequencies from the raw voltage data outside of 70 – 110 Hz, via a Butterworth band-pass filter. This specific frequency range has previously been associated with head-twitch events [36]. The data were transformed to absolute values, and local maxima were extracted to produce unipolar peaks. Finally, the conditional detection of new maxima from the previously extracted local maxima allowed for individual head-twitch events to be identified when (1) the peak prominence of the event exceeded the amplitude threshold (V), (2) the event was at least 200ms removed from any other event, and (3) the duration of the event did not exceed 90ms. The amplitude threshold (V) was calculated for each animal and, as previously validated [40], set to 15 standard deviations of the raw voltage data normalized to a mean of zero. Investigators were blinded to genotype and DOI dose until after completion of data analysis.

### Optogenetic light delivery

Optogenetic light, either yellow (590nm) for Ai39^+^ mice or blue (470nm) for Ai32^+^ mice, was delivered bilaterally using a bifurcated fiber bundle patch cord (ThorLabs, Cat. #: BFYL1LS01) attached to each fiber optic cannula via ceramic mating sleeves (RWD Life Science, 1.25mm). Fiber-coupled LEDs (ThorLabs, Cat. #: M590F3 yellow light, Cat #: MF4703 blue light) controlled by a LED driver (ThorLabs, Cat. #: LEDD1B) were used as light sources. Yellow light intensity was set to ∼ 0.26mW, and blue light intensity was set to ∼ 0.31mW for all HTR experiments. All light delivery was triggered by a Master-8 Pulse Stimulator (AMPI). During HTR experiments, Ai39^+^ mice received continuous yellow light, and Ai32^+^ mice received 5ms pulses of blue light delivered at 10 Hz, 5sec on/5sec off. Optogenetic light was delivered to both Ai39^+^ and Ai32^+^ mice immediately after drug injection for the full 30 min HTR recording session. “Light off” indicates that all components of the optogenetic light delivery system were connected and patch cords were attached to each mouse, but optogenetic light was not turned on. HTR experiments were completed for all genotypes under both light off and light on conditions, for each DOI dose tested.

### Statistical analysis

All statistical analyses were carried out using GraphPad Prism version 10 with significance set at *p*LJ<LJ0.05. Repeated measures two-way analysis of variance (ANOVA) with Šídák’s *post-hoc* tests were performed when the data were balanced (ie. equal *n* in all groups) and for conditions when multiple measures of the same subject were taken under different conditions and/or over time. When the data were unbalanced (ie. unequal *n* across groups), repeated measures, mixed effects analyses with Šídák’s *post-hoc* tests were performed. Geisser-Greenhouse correction (no assumption of sphericity) was applied for analyses in which the repeated measures factor had more than two levels. When there were at least three animals per sex, data were analyzed with sexes combined and with males and females separately. Data were plotted as meansLJ±LJstandard error of the mean (SEM).

## Results

To determine the location and role of 5-HT_2A_R-expressing neurons in the HTR to the psychedelic 5-HT_2A_R agonist DOI we used *Htr2a*-Cre mice [49,50]. We began by validating the location of *Htr2a*-Cre expression by crossing the line to the Ai14 Cre-dependent tdTomato reporter line of mice [51]. tdTomato expression is Cre-dependent and thus limited to 5-HT_2A_R expressing cells (Fig. 1). Analysis of tdTomato reporter expression in coronal and sagittal sections from *Htr2a*-Cre^+/-^;Ai14^+/-^ mice showed a similar anterior-posterior gradient to that of *in situ* hybridization, radioligand binding, and antibody staining studies in rodents [45–48]. High levels of tdTomato fluorescence are seen in the primary and secondary somatomotor areas, particularly in layers 2/3 and 5, the superficial layers of the dorsal and ventral anterior cingulate cortex, and in more lateral areas such as layer 5 of the visceral area (Fig. 1b). Also, as seen in our prior *Htr2a in situ* hybridization studies in mice [45], and previously published mRNA studies in rats [46,47], *Htr2a*-dependent Td Tomato reporter expression is notably absent from layers 2/3 and 4 of the primary and supplemental somatosensory areas. Outside of the cortex, low levels of 5-HT_2A_R protein are also seen in the caudate putamen, hypothalamus, piriform area, and olfactory tubercle, as well as in the medial and lateral septal nuclei.

**Figure 1.**
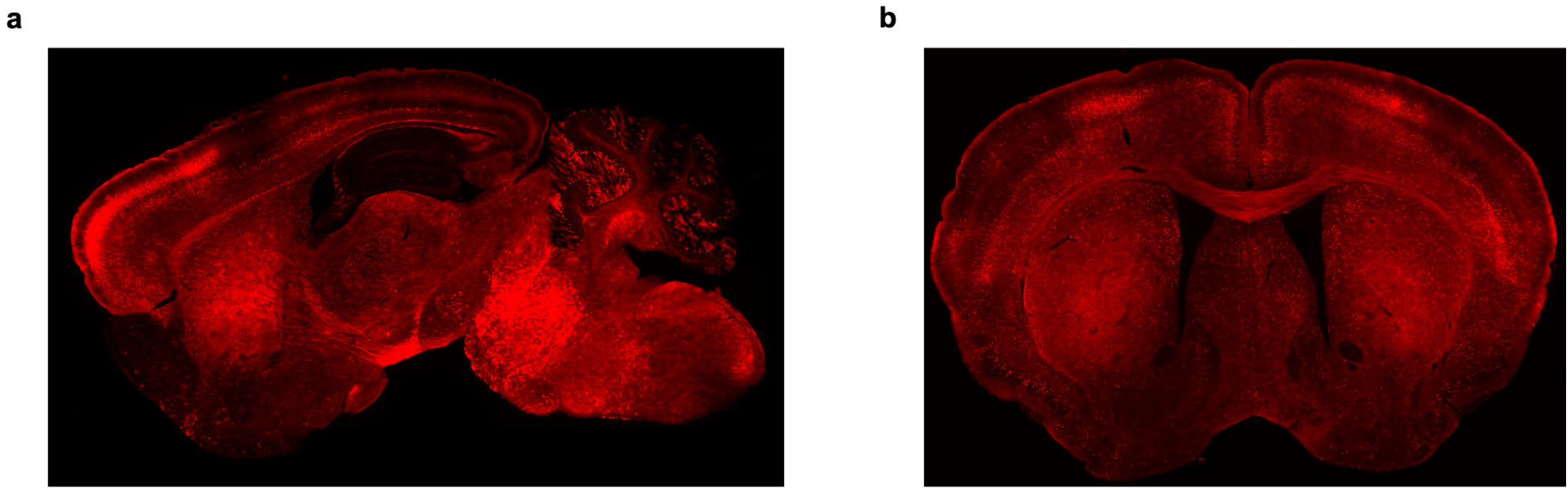
Fluorescent reporter distribution in *Htr2a* expressing neurons. *Htr2a-*Cre driven expression of Cre-dependent tdTomato parallels prior *in situ* hybridization and protein studies showing an antero-posterior gradient of cortical 5-HT_2A_R expression. Representative sagittal image (a) lateral 2.00mm and coronal image (b) bregma 1.10mm from *Htr2a*-Cre^+/-^;Ai14^+/-^ mice.

To manipulate the activity of 5-HT_2A_R-expressing neurons in the PFC, we crossed *Htr2a*-Cre mice to mice expressing Cre-dependent inhibitory and excitatory opsins. Since both halorhodopsin (HR) and channelrhodopsin (ChR) are expressed in a Cre-dependent manner, only 5-HT_2A_R+ neurons will be affected by optogenetic light delivery, allowing for cell-type specific modulation of neuronal activity (Fig. 2a). To validate the functional expression of HR and ChR, we performed *ex vivo* whole-cell patch clamp studies on PFC 5-HT_2A_R+ (EYFP expressing) neurons from *Htr2a*-Cre^+/-^;Ai39^+/-^ mice and *Htr2a*-Cre^+/-^;Ai32^+/-^ mice, respectively (Fig. 2b-e). We found that 100pA of current applied over 1sec to a 5-HT_2A_R+ PFC neuron from a *Htr2a*-Cre^+/-^;Ai39^+/-^ mouse cortical slice produced the expected depolarization and subsequent action potentials (Fig. 2c). When yellow light (590nm) and current (100pA) were applied simultaneously to the same neuron, the neuron depolarized but did fire any action potentials (Fig. 2c). Furthermore, application of yellow light in the absence of current prompted neuronal hyperpolarization (Fig. 2c). Exposure of 5-HT_2A_R+ PFC neurons from *Htr2a*-Cre^+/-^;Ai32^+/-^ mice to 1sec of continuous blue light (470nm) resulted in tetanic neuronal depolarization, while 5ms of 5Hz, 10Hz, or 20Hz pulses of blue light produced time locked depolarizations at the same frequencies (Fig. 2e).

**Figure 2.**
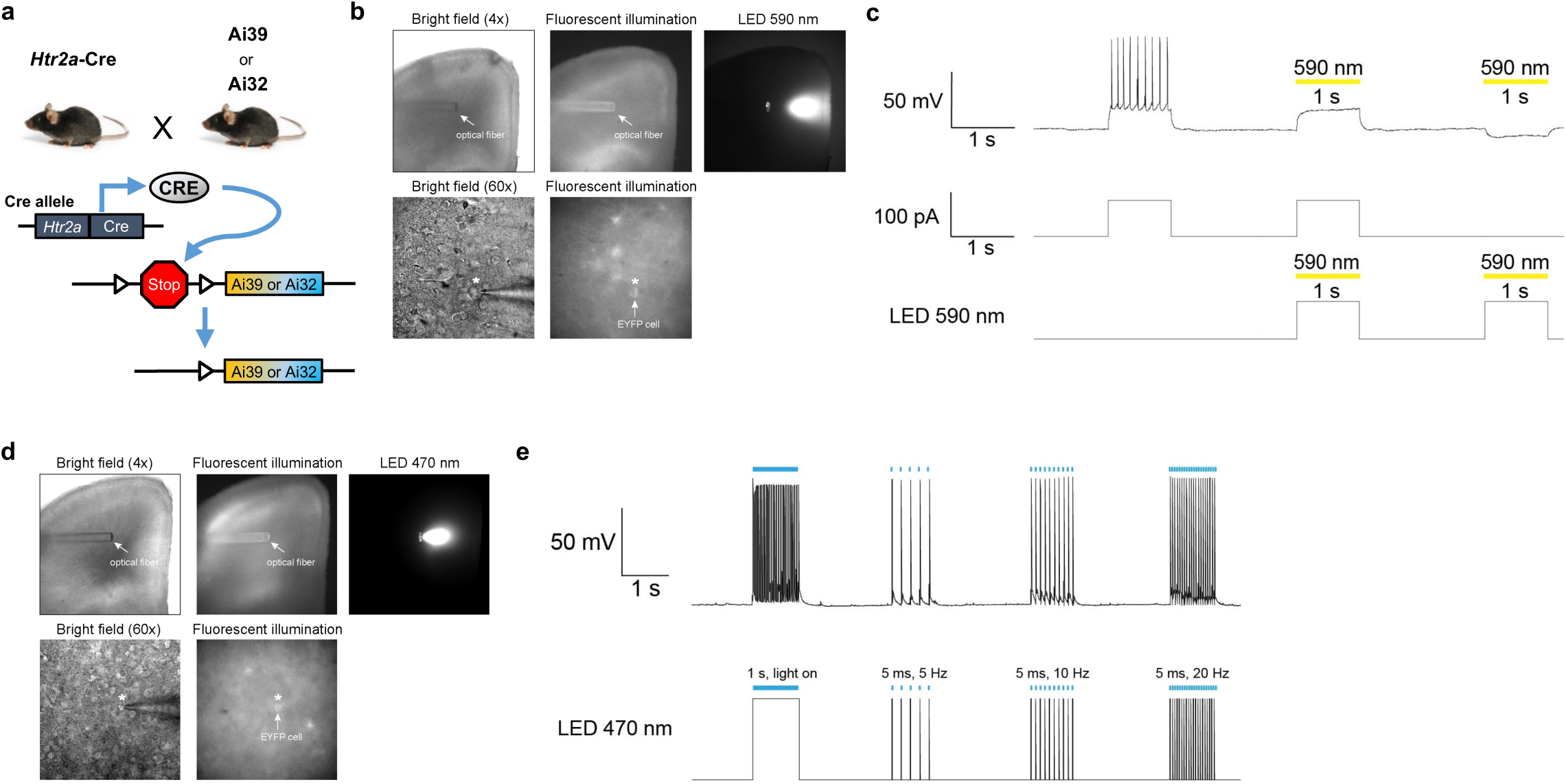
Whole-cell patch clamp validates functional expression of halorhodopsin and channelrhodopsin in cortical brain slices. (a) Schematic showing generation of mice expressing halorhodopsin (HR, Ai39) or channelrhodopsin (ChR, Ai32) in 5-HT_2A_R expressing neurons. (b) (Top) Images (4X) showing the position of the optical fiber on the PFC-containing slice. (Bottom) Images (60X) showing a patched PFC neuron expressing EYPF. (c) Voltage responses of EYFP positive PFC neuron to 1 s-long yellow light pulses (590 nm). (d) (Top) Images (4X) showing the position of the optical fiber on the PFC-containing slice. (Bottom) Images (60X) showing a patched PFC neuron expressing EYPF. **(e)** Voltage responses of EYFP positive PFC neuron to repetitive trains of 1 s-long or 5 ms-long blue light pulses (470 nm) at different frequencies (5Hz, 10 Hz and 20 Hz).

To determine if optogenetic inhibition of 5-HT_2A_R expressing neurons in the PFC prevents induction of HTRs in mice following treatment with the selective 5-HT_2A_R agonist DOI, we implanted bilateral fiber optic cannulae into the PFC to deliver yellow light (Fig. 3a). DOI administration (1mg/kg) significantly increased the number of HTRs in control mice (non-opsin expressing) compared to vehicle treated mice (Fig. 3b). However, optogenetic inhibition of 5-HT_2A_R+ cells in the PFC, in *Htr2a*-Cre^+/-^;Ai39^+/-^ (HR expressing) mice, prevented this DOI-induced increase in HTR (Fig. 3b), demonstrating that the activity of 5-HT_2A_R+ neurons in the PFC is necessary for the induction of HTR by the agonist DOI.

**Figure 3.**
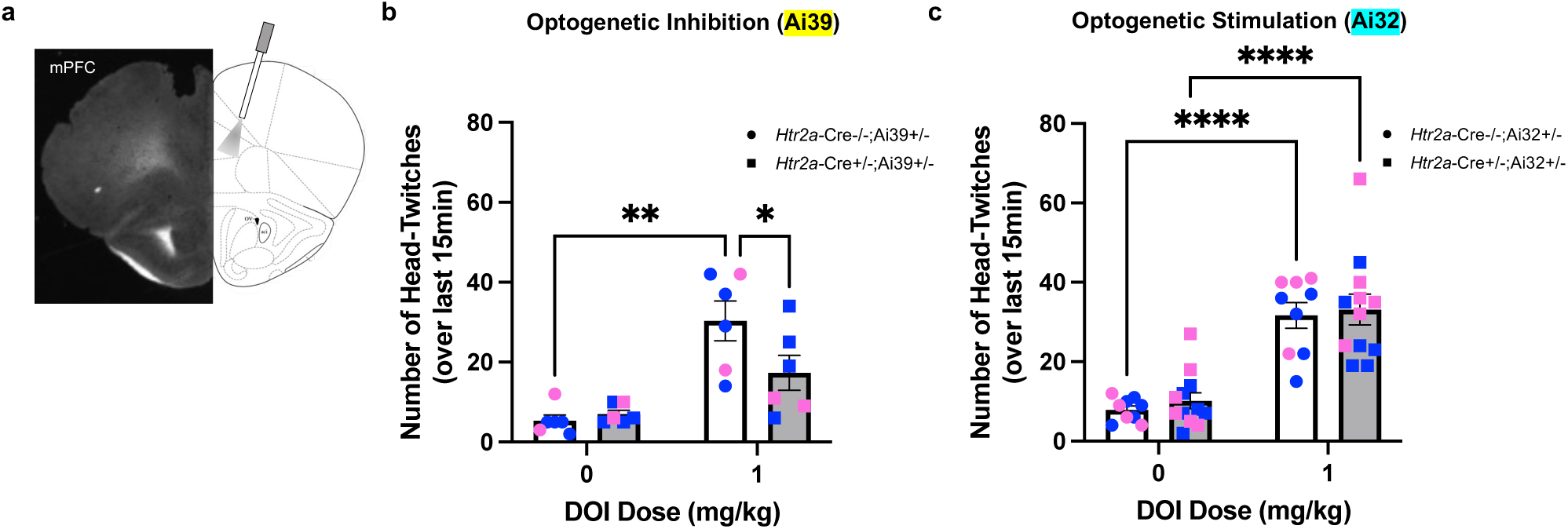
Activity of 5-HT_2A_R expressing neurons in the PFC is necessary, but not sufficient, to regulate DOI-induced HTRs. (a) Image of representative coronal brain section showing cannula track and atlas schematic showing location of bilateral optical fiber implantation in the mPFC. (b) Optogenetic inhibition (Ai39, yellow light) of 5-HT_2A_R expressing neurons prevents the increase in HTR by 1 mg/kg DOI, significant main effect of DOI (Two-way RM ANOVA, *p* = 0.0012), no effect of genotype (Two-way RM ANOVA, *p* = 0.0723). Post hoc analyses show significantly fewer HTRs in mice expressing HR in 5-HT_2A_R+ neurons compared to non-HR expressing mice. Optogenetic stimulation (blue light) does not induce HTR in the absence of DOI in mice expressing ChR in 5-HT_2A_R+ neurons (Ai32) compared with control (ChR non-expressing) mice. DOI induces a similar number of HTRs in both ChR expressing and non-expressing mice, significant main effect of DOI (Two-way RM ANOVA, *p* < 0.0001), no effect of genotype (Two-way RM ANOVA, *p* = 0.5073). Values represent means ± SEM. Post-hoc analyses corrected for multiple comparisons: * *p* < 0.05, ** *p* < 0.01, **** *p* < 0.0001, RM – repeated measures, blue – male, pink – female.

We next used ChR expressing mice to examine whether optogenetic simulation of 5-HT_2A_R positive neurons in the PFC was sufficient to induce HTRs in the absence of the agonist DOI. In vehicle treated mice, optogenetic stimulation of 5-HT_2A_R+ cells in the PFC did not induce HTRs (Fig. 3c), indicating that activation of PFC 5-HT_2A_R+ neurons is not sufficient to induce HTR in the absence of a 5-HT_2A_R agonist. Treatment with the agonist DOI (1mg/kg) increased the number of HTRs in both control and ChR-expressing mice when compared to vehicle treated mice (Fig. 3c). Notably, optogenetic simulation did not increase the HTR to DOI (1mg/kg) in ChR-expressing mice compared to non-opsin expressing mice (Fig. 3c).

These results suggest that optogenetic stimulation of PFC 5-HT_2A_R+ cells may not augment the HTR to DOI. However, it is possible a 1mg/kg dose of DOI may induce a maximum HTR in mice, thus representing a ceiling effect on this behavior. To address this question, we conducted additional studies with increased sample sizes, sufficient to analyze sexes independently, added optogenetic ‘lights off’ control conditions, and dose response studies to determine if stimulation of 5-HT_2A_R-expressing neurons in the PFC may augment the effect of lower doses of DOI.

To determine whether the genotype of *Htr2a*-Cre;Ai39 alone, in the absence of optogenetic inhibition (yellow light), may affect HTR behavior, we compared the effect DOI (1mg/kg) on HTRs in control mice and mice expressing HR in 5-HT_2A_R+ cells (male and female data combined, Fig. 4a). Figure 4a shows that DOI increased HTR in both groups and that there was no significant difference in the number of HTRs between mice expressing HR in 5-HT_2A_R+ cells (*Htr2a*-Cre^+/-^;Ai39^+/-^) and non-opsin expressing control mice in response to either vehicle (0mg/kg) or DOI (1mg/kg) (males and females combined). The same result was seen under lights-off conditions when males and females were analyzed separately (Fig. 4b and 4c, respectively). These results indicated that in the absence of optogenetic inhibition the *Htr2a*-Cre;Ai39 genotype alone does not affect HTR behavior at baseline or in response to DOI (1mg/kg).

**Figure 4.**
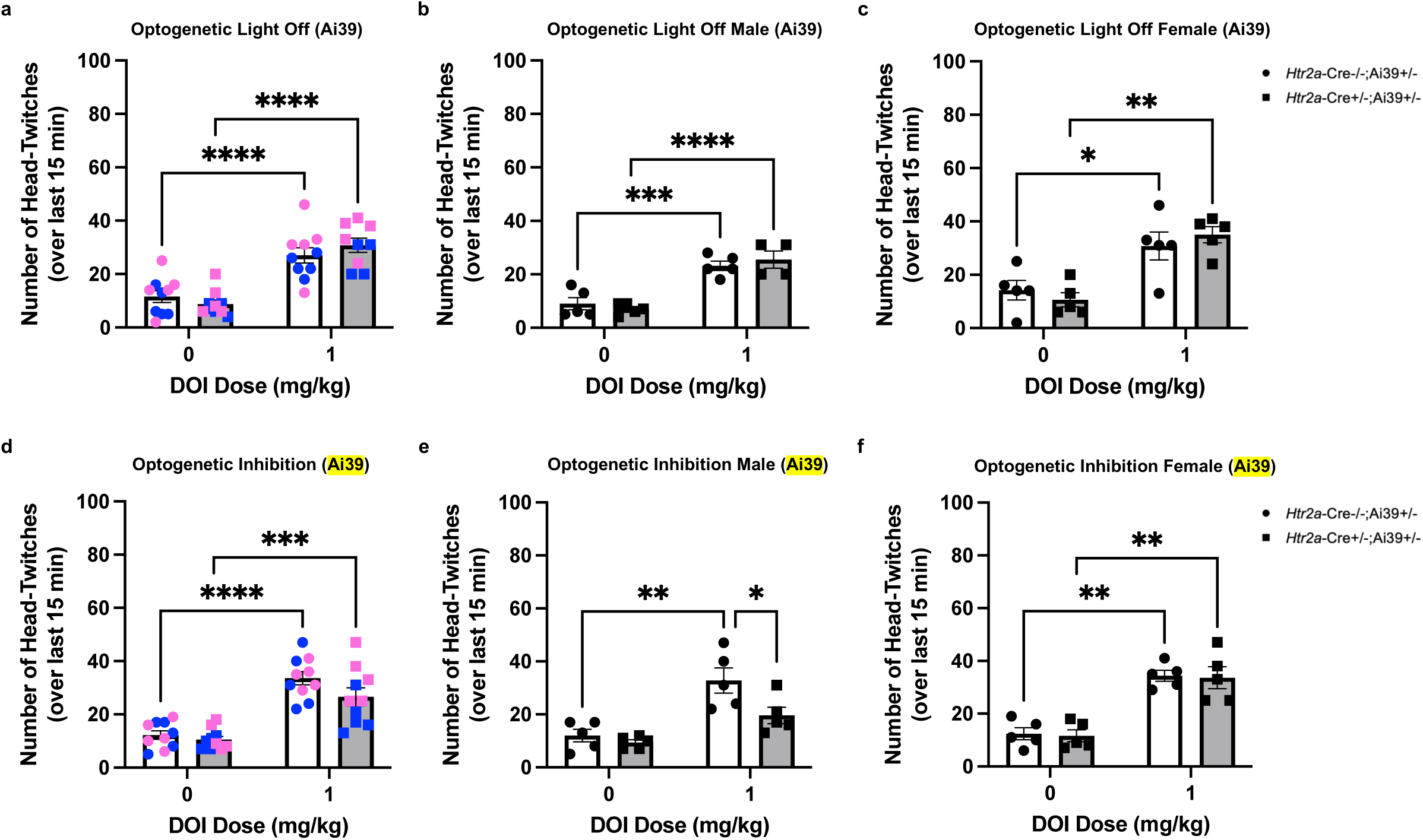
Optogenetic inhibition of PFC 5-HT_2A_R expressing neurons prevents induction of HTR to DOI in male mice. (a) Under lights-off conditions, when male and female mice are analyzed together, DOI-induces HTR similarly in mice expressing HR in 5-HT_2A_R+ neurons of the PFC and control (HR non-expressing) mice, significant main effect of DOI (Mixed-effects analysis, RM, *p* < 0.0001) and no effect of genotype (Mixed-effects analysis, RM, *p* = 0.8373). When each sex was analyzed separately, under lights-off conditions there were also no differences between genotypes in the HTR to DOI in (b) males, significant main effect of DOI (Mixed-effects analysis, RM, *p* < 0.0001) and no effect of genotype (Mixed-effects analysis, RM, *p* = 0.9435), or (c) females, significant main effect of DOI (Two-way RM ANOVA, *p* = 0.0012) and no effect of genotype (Two-way RM ANOVA, *p* = 0.9306). (d) In the presence of optogenetic inhibition, when male and female mice are analyzed together, DOI-induces HTR similarly in mice expressing HR in 5-HT_2A_R+ neurons of the PFC and control (HR non-expressing) mice, significant main effect of DOI (Two-way RM ANOVA, *p* < 0.0001) and no effect of genotype (Two-way RM ANOVA, *p* = 0.0924). (e) In males, inhibition of 5-HT_2A_R expressing neurons in the PFC prevented DOI-induced HTR, significant main effect of DOI (Two-way RM ANOVA, *p* = 0.0004) and no effect of genotype (Two-way RM ANOVA, *p* = 0.0568). Post-hoc analyses showed significantly fewer HTRs in male mice expressing HR in 5-HT_2A_R+ neurons compared to non-HR expressing male mice. (f) In females, inhibition of 5-HT_2A_R expressing neurons in the PFC did not prevent DOI-induced HTR, significant main effect of DOI (Two-way RM ANOVA, *p* = 0.0001), no effect of genotype (Two-way RM ANOVA, *p* = 0.7527). Values represent means ± SEM. Post-hoc analyses adjusted for multiple comparisons: * *p* < 0.05, ** *p* < 0.01, *** *p* < 0.001, **** *p* < 0.0001, RM – repeated measures, blue – male, pink – female.

Next, we tested if optogenetic inhibition of 5-HT_2A_R+ neurons in the PFC altered DOI-induced HTR in male and female HR-expressing mice compared to male and female non-HR expressing control mice. When both sexes were analyzed together, optogenetic inhibition did not alter DOI-induced HTR in mice that express HR in 5-HT_2A_R+ neurons compared to HR non-expressing mice (Fig. 4d). However, when testing males and females separately we identified sex-dependent differences. In *male* mice, under continuous optogenetic inhibition, control non HR-expressing mice showed the expected increase in HTR to DOI (1mg/kg) compared to vehicle treatment (Fig. 4e). In contrast, *male* mice expressing HR in 5-HT_2A_R+ neurons, DOI (1mg/kg) administration failed to significantly increase HTRs in the presence of yellow light (Fig. 4e). And following DOI, male mice expressing HR showed significantly fewer HTRs than HR non-expressing male mice (Fig. 4e). These findings replicated the results of our initial study (Fig. 3b). In contrast, in *female* mice under continuous optogenetic inhibition, control non HR-expressing *and* HR-expressing mice showed significant increases in HTR to DOI (1mg/kg) compared to vehicle treatment (Fig. 4f). Additionally, following DOI, female control non HR-expressing and HR-expressing mice both showed a similar number of HTRs (Fig. 4e). Together, these results show that optogenetic inhibition of PFC 5-HT_2A_R expressing neurons prevents induction of HTR to DOI in male mice but not female mice.

Before testing the effect of optogenetic stimulation of 5-HT_2A_R expressing neurons on the HTR to lower doses of DOI, we wanted to ensure that there was no effect of *Htr2a*-Cre;Ai32 genotype alone. To address this, we compared the effect multiple DOI doses (0.1mg/kg, 0.3mg/kg, and 1mg/kg) on HTRs in control mice and mice expressing ChR in 5-HT_2A_R+ cells. In the absence of optogenetic simulation, DOI induced HTR similarly in ChR-expressing mice and non-ChR expressing mice when both sexes were analyzed together (Fig. 5a). In males, without optogenetic stimulation (blue light), DOI induced HTR was similar in ChR-expressing and non-expressing male mice (Fig. 5b). This same effect was seen in female ChR-expressing and non-ChR expressing mice in the absence of optogenetic simulation (Fig. 5c).

**Figure 5.**
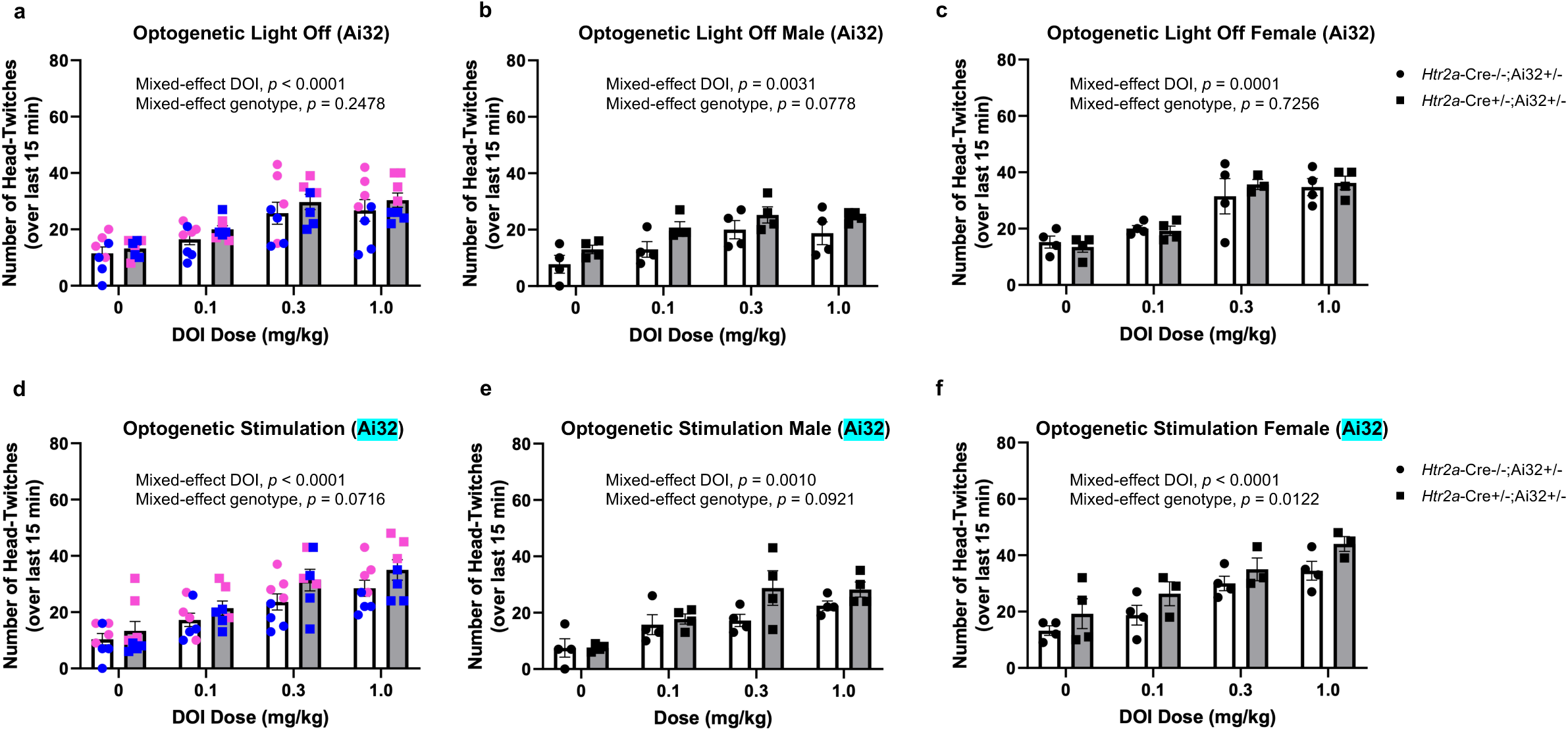
Optogenetic stimulation of PFC 5-HT_2A_R expressing neurons augments the induction of HTR to DOI in female mice. (a) In the absence of optogenetic stimulation (i.e., light-off conditions), there was not difference in the dose dependent effect of DOI on HTR between genotypes when both sexes were analyzed together, significant main effect of DOI (Mixed-effects analysis, RM, Geisser-Greenhouse correction, *p* < 0.0001) and no effect of genotype (Mixed-effects analysis, RM, Geisser-Greenhouse correction, *p* = 0.2478), or separately (b) in males, significant main effect of DOI (Mixed-effects analysis, RM, Geisser-Greenhouse correction, *p* = 0.0031) and no effect of genotype (Mixed-effects analysis, RM, Geisser-Greenhouse correction, *p* = 0.0778) and (c) in females, significant main effect of DOI (Mixed-effects analysis, RM, Geisser-Greenhouse correction, *p* = 0.0001) and no effect of genotype (Mixed-effects analysis, RM, Geisser-Greenhouse correction, *p* = 0.7256). (d) In the presence of blue light, when both sexes are analyzed together, stimulation of 5-HT_2A_R expressing neurons in the PFC did not alter DOI-induced HTR, no effect of genotype (Mixed-effects analysis, RM, Geisser-Greenhouse correction, *p* = 0.0716) and significant main effect of DOI (Mixed-effects analysis, RM, Geisser-Greenhouse correction, *p* < 0.0001). (e) Similarly, in males, stimulation of of 5-HT_2A_R expressing neurons in the PFC did not alter DOI-induced HTR, no effect of genotype (Mixed-effects analysis, RM, Geisser-Greenhouse correction, *p* = 0.0921) and significant main effect of DOI (Mixed-effects analysis, RM, Geisser-Greenhouse correction, *p* = 0.0010). (f) However, in females, stimulation of 5-HT_2A_R expressing neurons in the PFC increases the number of DOI-induced HTRs compared to non-ChR expressing controls, significant main effect of genotype (Mixed-effects analysis, RM, Geisser-Greenhouse correction, *p* = 0.0122) and significant effect of DOI (Mixed-effects analysis, RM, Geisser-Greenhouse correction, *p* < 0.0001). Values represent means ± SEM. No post-hoc analyses were significant. RM – repeated measures, blue – male, pink – female.

We then examined whether optogenetic stimulation of 5-HT_2A_R positive neurons in the PFC could augment the HTR of mice to intermediate doses of DOI (Fig. 5). Under optogenetic simulation conditions, treatment with DOI induced HTR similarly in ChR-expressing mice and non-ChR expressing controls when both sexes were analyzed together (Fig. 5d). Similarly, in *males* alone, optogenetic stimulation of 5-HT_2A_R+ neurons of the PFC did not alter DOI-induced HTR (Fig. 5e). Interestingly, in *female* mice, optogenetic stimulation increased the number of DOI-induced HTRs in ChR-expressing *female* mice compared to non-ChR expressing *female* mice (Fig. 5f). In addition, as we had seen in the first study (Fig. 3c), for both male and female mice, optogenetic stimulation of 5-HT_2A_R+ neurons in the PFC was not sufficient to induce HTRs in vehicle-treated ChR-expressing mice in comparison to non-ChR expressing mice (Fig. 5e, 5f).

## Discussion

The results we report here provide novel insight into the brain region and role of 5-HT_2A_R+ neuronal activity in regulation of the HTR to the agonist DOI. Our findings demonstrate that optogenetic activation and inhibition of PFC 5-HT_2A_R+ neurons can augment the induction of HTRs in mice by DOI in a sex-specific manner. We show that activation of PFC 5-HT_2A_R+ neurons is not sufficient to induce HTRs in the absence of the 5-HT_2A_R agonist DOI, regardless of sex. Furthermore, we show that activation of 5-HT_2A_R+ neurons in the PFC is *necessary* for induction of HTRs in *male* mice, but may not be in *female* mice. Finally, our results also demonstrate that, in *female* mice, activation of 5-HT_2A_R+ neurons in the PFC augments the number of HTRs elicited by various doses of DOI, a result absent in *male* mice.

Our findings that optogenetic inhibition and activation of 5-HT_2A_R expressing neurons in the PFC can alter HTR of mice to the agonist DOI provide further validation that the PFC is a region important in the regulation of the HTR behavior. Willins and Meltzer found that direct injection of 5-HT_2A_R agonists into the medial prefrontal cortex (mPFC) induces HTR in rats [38]. Specifically, bilateral injection of the selective 5-HT_2A_R agonist DOI and the 5-HT_2A_R partial agonist m-chloro-phenylpiperazine (mCPP) into the mPFC of male rats elicits a dose-dependent induction of HTR that is attenuated by pretreatment with either of the 5-HT_2A_R antagonists ketanserin or MDL 100,907 [38]. Although some studies have suggested that the frontal cortex is not involved in the HTR behavior [56,57], the findings of Willins and Meltzer, together with the results presented here, suggest that 1) HTR is mediated by the selective activation of 5-HT_2A_Rs and 2) the activity of cortical 5-HT_2A_R expressing neurons is required, in a sex dependent manner, for psychedelic-induced HTRs in rodents [38].

One of the interesting findings we report here are the sex differences in the effects of activation and inhibition of PFC 5-HT_2A_R+ neurons in response to DOI-induced HTR. Few prior studies have examined the impact of sex on HTR. Darmani and colleagues reported that HTRs elicited by DOI did not differ between male and female mice [58]. An earlier study by Boulton and Handley showed that estrus cycle had no effect on HTR following systemic administration (i.p. injection) of the amino acid serotonin precursor 5-hydroxytryptophan (5-HTP) but that female mice (combined estrus and diestrus) had significantly fewer HTRs in response to 5-HTP than male mice [59].

More recent studies exploring the impact of sex on HTR have shown significantly more HTRs in female mice than in male mice when number of HTRs were tracked using time course assays in both male and female C57BL/6J mice following administration of DOI [60]. These sex differences appeared to be strain specific as 129S6/SvEv mice did not show a sex difference in this response [60]. The number of spontaneous HTRs (i.e., in the absence of vehicle or drug injection), and HTRs following vehicle administration, were not different between sexes in either the C57BL/6J or 129S6/SvEv strains [60]. Notably, this sex difference in the C57BL/6J mice appears to be dose dependent, as 0.5mg/kg of DOI resulted in a similar number if HTRs in males and females, but HTRs induced by 2.0mg/kg of DOI appeared to elicit a greater number of HTRs in female compared to male mice (reviewed in Fig. 1a of [60]). Jaster and colleagues went on to propose that sex differences in DOI-induced HTR in C57BL/6J mice may be explained by different pharmacokinetics of the drug in male and female mice. They found that frontal cortex and plasma concentration of DOI were lower in female compared to male C57BL/6J mice at 30min and 60min after 2mg/kg i.p. injection of DOI [60].

Previous rat studies have shown that *Htr2a* mRNA expression was comparable between males and females in all brain regions except for the ventromedial hypothalamic nuclei, in which female rat has lower levels of *Htr2a* expression [61]. Yet in the same study, binding of 5-HT_2A_Rs in female rats, measured via [^3^H]ketanserin, was significantly higher in all regions of the hippocampus compared to males [61]. Soloff and colleagues conducted a PET study in humans using [^18^F]altanserin to assess sex differences in 5-HT_2A_R binding throughout the forebrain in healthy male and female subjects. Notably, [^18^F]altanserin binding was greater in males, compared to females, in the following brain regions: hippocampus, lateral orbital frontal cortex, left medial frontal cortex [62]. These sex specific DOI pharmacokinetics and receptor density variations may provide a potential explanation for our sex-specific findings.

Our prior work has demonstrated that 5-HT_2A_R levels in the frontal cortex change dramatically in response to experience and events [45,63]. In addition, a significant body of research has shown that levels of 5-HT_2A_R mRNA and protein, as well as 5-HT_2A_R mediated behaviors like HTR, are elevated in response to various environmental stimuli such as, but not limited to, physical immobilization, toe pinch, and social stress (reviewed in [64]). Given this dynamic nature of 5-HT_2A_Rs, it is possible that response to agonists may be affected by environmental events. This is particularly intriguing in the context of psychedelics, as setting plays an important role in the psychedelic experience [65], suggesting another way in which the environment is crucial for the response to psychedelics.

The surge of recent studies demonstrating compelling therapeutic effects of psychedelics underscores the importance of elucidating the molecular mechanisms underlying these powerful drugs. Recently, many clinical trials have reported that one or two doses of psilocybin can produce substantial and rapid antidepressant effects [18,25–29]. Furthermore, there is some evidence that just two doses of psilocybin can result in a significant reduction of depressive symptoms that lasts for at least 6 months [18,25]. Although limited by small sample size, and in one case open label [18], if these results hold true in larger clinical trials, these findings could transform treatment for depression, for which second-generation antidepressant medications often take 4-6 weeks to achieve similar effects.

Together, our findings suggest that activity of 5-HT_2A_R-expressing neurons in the PFC is not sufficient for HTR to psychedelic drugs in mice but may be necessary in a sex specific manner. Since HTR is highly correlated with the perceptual effects of psychedelic compounds, our results provide support that the prefrontal cortex may be a region involved in the therapeutic effects of these drugs. These findings thereby begin to tease apart a possible mechanism through which psychedelic drugs elicit therapeutic effects in humans and thus may contribute to the treatment of neuropsychiatric illness.

## Acknowledgements

We are grateful to K. Meyers, PhD for assistance with the original building of the automated head-twitch response apparatus and for providing instruction for experimental protocols and to Mario de la Fuente Revenga, PhD for expert advice on MatLab programming. Funding: This work was supported by National Institutes of Health R01 MH097803 (ALG and SQ).

## Conflicts of Interest

The authors have no conflicts of interest to report.

## Contributions

ABO: Investigation, Formal analysis, Visualization, Writing – original draft, Writing – review & editing. JW: Investigation, Formal analysis, Visualization. JC: Investigation, Resources. CH: Formal analysis, SQ: Conceptualization, Funding acquisition, Supervision, Resources, Writing – review & editing. ALG: Conceptualization, Funding acquisition, Supervision, Writing – review & editing.

## Notes

### Competing Interest Statement

The authors have declared no competing interest.

## References

1 Nichols DE. Hallucinogens. Pharmacol Ther. 2004;101(2):131–81.

2 Halberstadt AL, Geyer MA. Multiple receptors contribute to the behavioral effects of indoleamine hallucinogens. Neuropharmacology. 2011;61(3):364–81.

3 Kaye WH, Frank GK, Bailer UF, Henry SE, Meltzer CC, Price JC, et al. Serotonin alterations in anorexia and bulimia nervosa: new insights from imaging studies. Physiol Behav. 2005;85(1):73–81.

4 Spigset O, Andersen T, Hagg S, Mjondal T. Enhanced platelet serotonin 5-HT2A receptor binding in anorexia nervosa and bulimia nervosa. Eur Neuropsychopharmacol. 1999;9(6):469–73.

5 Audenaert K, Van Laere K, Dumont F, Vervaet M, Goethals I, Slegers G, et al. Decreased 5-HT2a receptor binding in patients with anorexia nervosa. J Nucl Med. 2003;44(2):163–9.

6 Selvaraj S, Arnone D, Cappai A, Howes O. Alterations in the serotonin system in schizophrenia: a systematic review and meta-analysis of postmortem and molecular imaging studies. Neurosci Biobehav Rev. 2014;45:233–45.

7 Dean B, Hayes W. Decreased frontal cortical serotonin2A receptors in schizophrenia. Schizophr Res. 1996;21(3):133–9.

8 Rasmussen H, Frokjaer VG, Hilker RW, Madsen J, Anhoj S, Oranje B, et al. Low frontal serotonin 2A receptor binding is a state marker for schizophrenia? Eur Neuropsychopharmacol. 2016;26(7):1248–50.

9 Hurlemann R, Matusch A, Kuhn KU, Berning J, Elmenhorst D, Winz O, et al. 5-HT2A receptor density is decreased in the at-risk mental state. Psychopharmacology (Berl). 2008;195(4):579–90.

10 Ngan ET, Yatham LN, Ruth TJ, Liddle PF. Decreased serotonin 2A receptor densities in neuroleptic-naive patients with schizophrenia: A PET study using [(18)F]setoperone. Am J Psychiatry. 2000;157(6):1016–8.

11 Lopez-Figueroa AL, Norton CS, Lopez-Figueroa MO, Armellini-Dodel D, Burke S, Akil H, et al. Serotonin 5-HT1A, 5-HT1B, and 5-HT2A receptor mRNA expression in subjects with major depression, bipolar disorder, and schizophrenia. Biol Psychiatry. 2004;55(3):225–33.

12 Matsumoto I, Inoue Y, Iwazaki T, Pavey G, Dean B. 5-HT2A and muscarinic receptors in schizophrenia: a postmortem study. Neurosci Lett. 2005;379(3):164–8.

13 Serretti A, Drago A, De Ronchi D. HTR2A gene variants and psychiatric disorders: a review of current literature and selection of SNPs for future studies. Curr Med Chem. 2007;14(19):2053–69.

14 Burnet PW, Eastwood SL, Harrison PJ. 5-HT1A and 5-HT2A receptor mRNAs and binding site densities are differentially altered in schizophrenia. Neuropsychopharmacology. 1996;15(5):442–55.

15 Celada P, Puig M, Amargos-Bosch M, Adell A, Artigas F. The therapeutic role of 5-HT1A and 5-HT2A receptors in depression. J Psychiatry Neurosci. 2004;29(4):252–65.

16 Bhagwagar Z, Hinz R, Taylor M, Fancy S, Cowen P, Grasby P. Increased 5-HT(2A) receptor binding in euthymic, medication-free patients recovered from depression: a positron emission study with [(11)C]MDL 100,907. Am J Psychiatry. 2006;163(9):1580–7.

17 Petit AC, Quesseveur G, Gressier F, Colle R, David DJ, Gardier AM, et al. Converging translational evidence for the involvement of the serotonin 2A receptor gene in major depressive disorder. Prog Neuropsychopharmacol Biol Psychiatry. 2014;54:76–82.

18 Carhart-Harris RL, Bolstridge M, Day CMJ, Rucker J, Watts R, Erritzoe DE, et al. Psilocybin with psychological support for treatment-resistant depression: six-month follow-up. Psychopharmacology (Berl). 2018;235(2):399–408.

19 Bogenschutz MP, Ross S, Bhatt S, Baron T, Forcehimes AA, Laska E, et al. Percentage of Heavy Drinking Days Following Psilocybin-Assisted Psychotherapy vs Placebo in the Treatment of Adult Patients With Alcohol Use Disorder: A Randomized Clinical Trial. JAMA Psychiatry. 2022;79(10):953–62.

20 Johnson MW, Garcia-Romeu A, Cosimano MP, Griffiths RR. Pilot study of the 5-HT2AR agonist psilocybin in the treatment of tobacco addiction. J Psychopharmacol. 2014;28(11):983–92.

21 Johnson MW, Garcia-Romeu A, Griffiths RR. Long-term follow-up of psilocybin-facilitated smoking cessation. Am J Drug Alcohol Abuse. 2017;43(1):55–60.

22 Nichols DE, Johnson MW, Nichols CD. Psychedelics as Medicines: An Emerging New Paradigm. Clin Pharmacol Ther. 2017;101(2):209–19.

23 Holze F, Gasser P, Muller F, Dolder PC, Liechti ME. Lysergic Acid Diethylamide-Assisted Therapy in Patients With Anxiety With and Without a Life-Threatening Illness: A Randomized, Double-Blind, Placebo-Controlled Phase II Study. Biol Psychiatry. 2023;93(3):215–23.

24 Grob CS, Danforth AL, Chopra GS, Hagerty M, McKay CR, Halberstadt AL, et al. Pilot study of psilocybin treatment for anxiety in patients with advanced-stage cancer. Arch Gen Psychiatry. 2011;68(1):71–8.

25 Griffiths RR, Johnson MW, Carducci MA, Umbricht A, Richards WA, Richards BD, et al. Psilocybin produces substantial and sustained decreases in depression and anxiety in patients with life-threatening cancer: A randomized double-blind trial. J Psychopharmacol. 2016;30(12):1181–97.

26 Ross S, Bossis A, Guss J, Agin-Liebes G, Malone T, Cohen B, et al. Rapid and sustained symptom reduction following psilocybin treatment for anxiety and depression in patients with life-threatening cancer: a randomized controlled trial. J Psychopharmacol. 2016;30(12):1165–80.

27 Carhart-Harris RL, Bolstridge M, Rucker J, Day CM, Erritzoe D, Kaelen M, et al. Psilocybin with psychological support for treatment-resistant depression: an open-label feasibility study. Lancet Psychiatry. 2016;3(7):619–27.

28 Davis AK, Barrett FS, May DG, Cosimano MP, Sepeda ND, Johnson MW, et al. Effects of Psilocybin-Assisted Therapy on Major Depressive Disorder: A Randomized Clinical Trial. JAMA Psychiatry. 2021;78(5):481–89.

29 Goodwin GM, Aaronson ST, Alvarez O, Arden PC, Baker A, Bennett JC, et al. Single-Dose Psilocybin for a Treatment-Resistant Episode of Major Depression. N Engl J Med. 2022;387(18):1637–48.

30 Schreiber R, Brocco M, Audinot V, Gobert A, Veiga S, Millan MJ. (1-(2,5-dimethoxy-4 iodophenyl)-2-aminopropane)-induced head-twitches in the rat are mediated by 5-hydroxytryptamine (5-HT) 2A receptors: modulation by novel 5-HT2A/2C antagonists, D1 antagonists and 5-HT1A agonists. J Pharmacol Exp Ther. 1995;273(1):101–12.

31 Fantegrossi WE, Simoneau J, Cohen MS, Zimmerman SM, Henson CM, Rice KC, et al. Interaction of 5-HT2A and 5-HT2C receptors in R(-)-2,5-dimethoxy-4-iodoamphetamine-elicited head twitch behavior in mice. J Pharmacol Exp Ther. 2010;335(3):728–34.

32 Darmani NA, Martin BR, Pandey U, Glennon RA. Do functional relationships exist between 5-HT1A and 5-HT2 receptors? Pharmacol Biochem Behav. 1990;36(4):901–6.

33 Fox MA, Stein AR, French HT, Murphy DL. Functional interactions between 5-HT2A and presynaptic 5-HT1A receptor-based responses in mice genetically deficient in the serotonin 5-HT transporter (SERT). Br J Pharmacol. 2010;159(4):879–87.

34 Gonzalez-Maeso J, Yuen T, Ebersole BJ, Wurmbach E, Lira A, Zhou M, et al. Transcriptome fingerprints distinguish hallucinogenic and nonhallucinogenic 5-hydroxytryptamine 2A receptor agonist effects in mouse somatosensory cortex. J Neurosci. 2003;23(26):8836–43.

35 Gonzalez-Maeso J, Weisstaub NV, Zhou M, Chan P, Ivic L, Ang R, et al. Hallucinogens recruit specific cortical 5-HT(2A) receptor-mediated signaling pathways to affect behavior. Neuron. 2007;53(3):439–52.

36 Halberstadt AL, Geyer MA. Characterization of the head-twitch response induced by hallucinogens in mice: detection of the behavior based on the dynamics of head movement. Psychopharmacology (Berl). 2013;227(4):727–39.

37 Schmid CL, Raehal KM, Bohn LM. Agonist-directed signaling of the serotonin 2A receptor depends on beta-arrestin-2 interactions in vivo. Proc Natl Acad Sci U S A. 2008;105(3):1079–84.

38 Willins DL, Meltzer HY. Direct injection of 5-HT2A receptor agonists into the medial prefrontal cortex produces a head-twitch response in rats. J Pharmacol Exp Ther. 1997;282(2):699–706.

39 Glennon RA, Young R, Rosecrans JA. Antagonism of the effects of the hallucinogen DOM and the purported 5-HT agonist quipazine by 5-HT2 antagonists. Eur J Pharmacol. 1983;91(2-3):189–96.

40 de la Fuente Revenga M, Shin JM, Vohra HZ, Hideshima KS, Schneck M, Poklis JL, et al. Fully automated head-twitch detection system for the study of 5-HT(2A) receptor pharmacology in vivo. Sci Rep. 2019;9(1):14247.

41 Dinis-Oliveira RJ. Metabolism of psilocybin and psilocin: clinical and forensic toxicological relevance. Drug Metab Rev. 2017;49(1):84–91.

42 Saulin A, Savli M, Lanzenberger R. Serotonin and molecular neuroimaging in humans using PET. Amino Acids. 2012;42(6):2039–57.

43 Paterson LM, Kornum BR, Nutt DJ, Pike VW, Knudsen GM. 5-HT radioligands for human brain imaging with PET and SPECT. Med Res Rev. 2013;33(1):54–111.

44 Beliveau V, Ganz M, Feng L, Ozenne B, Hojgaard L, Fisher PM, et al. A High-Resolution In Vivo Atlas of the Human Brain’s Serotonin System. J Neurosci. 2017;37(1):120–28.

45 Zhao X, Ozols AB, Meyers KT, Campbell J, McBride A, Marballi KK, et al. Acute sleep deprivation upregulates serotonin 2A receptors in the frontal cortex of mice via the immediate early gene Egr3. Mol Psychiatry. 2022;27(3):1599–610.

46 Mengod G, Pompeiano M, Martinez-Mir MI, Palacios JM. Localization of the mRNA for the 5-HT2 receptor by in situ hybridization histochemistry. Correlation with the distribution of receptor sites. Brain Res. 1990;524(1):139–43.

47 Pompeiano M, Palacios JM, Mengod G. Distribution of the serotonin 5-HT2 receptor family mRNAs: comparison between 5-HT2A and 5-HT2C receptors. Brain Res Mol Brain Res. 1994;23(1-2):163–78.

48 Weber ET, Andrade R. Htr2a Gene and 5-HT(2A) Receptor Expression in the Cerebral Cortex Studied Using Genetically Modified Mice. Front Neurosci. 2010;4.

49 Gong S, Zheng C, Doughty ML, Losos K, Didkovsky N, Schambra UB, et al. A gene expression atlas of the central nervous system based on bacterial artificial chromosomes. Nature. 2003;425(6961):917–25.

50 Gong S, Doughty M, Harbaugh CR, Cummins A, Hatten ME, Heintz N, et al. Targeting Cre recombinase to specific neuron populations with bacterial artificial chromosome constructs. J Neurosci. 2007;27(37):9817–23.

51 Madisen L, Zwingman TA, Sunkin SM, Oh SW, Zariwala HA, Gu H, et al. A robust and high-throughput Cre reporting and characterization system for the whole mouse brain. Nat Neurosci. 2010;13(1):133–40.

52 Madisen L, Mao T, Koch H, Zhuo JM, Berenyi A, Fujisawa S, et al. A toolbox of Cre-dependent optogenetic transgenic mice for light-induced activation and silencing. Nat Neurosci. 2012;15(5):793–802.

53 Zeng H. Direct Database Submission 2011/06/21_Dl1gAf. MGI Direct Data Submission. 2011.

54 Lein ES, Hawrylycz MJ, Ao N, Ayres M, Bensinger A, Bernard A, et al. Genome-wide atlas of gene expression in the adult mouse brain. Nature. 2007;445(7124):168-76.

55 Allen Mouse Brain Atlas. 2004. mouse.brain-map.org.

56 Lucki I, Minugh-Purvis N. Serotonin-induced head shaking behavior in rats does not involve receptors located in the frontal cortex. Brain Res. 1987;420(2):403–6.

57 Bedard P, Pycock CJ. “Wet-dog” shake behaviour in the rat: a possible quantitative model of central 5-hydroxytryptamine activity. Neuropharmacology. 1977;16(10):663–70.

58 Darmani NA, Shaddy J, Gerdes CF. Differential ontogenesis of three DOI-induced behaviors in mice. Physiol Behav. 1996;60(6):1495–500.

59 Boulton CS, Handley SL. Factors modifying the head-twitch response to 5-hydroxytryptophan. Psychopharmacologia. 1973;31(3):205–14.

60 Jaster AM, Younkin J, Cuddy T, de la Fuente Revenga M, Poklis JL, Dozmorov MG, et al. Differences across sexes on head-twitch behavior and 5-HT(2A) receptor signaling in C57BL/6J mice. Neurosci Lett. 2022;788:136836.

61 Zhang L, Ma W, Barker JL, Rubinow DR. Sex differences in expression of serotonin receptors (subtypes 1A and 2A) in rat brain: a possible role of testosterone. Neuroscience. 1999;94(1):251–9.

62 Soloff PH, Price JC, Mason NS, Becker C, Meltzer CC. Gender, personality, and serotonin-2A receptor binding in healthy subjects. Psychiatry Res. 2010;181(1):77–84.

63 Maple AM, Zhao X, Elizalde DI, McBride AK, Gallitano AL. Htr2a Expression Responds Rapidly to Environmental Stimuli in an Egr3-Dependent Manner. ACS Chem Neurosci. 2015;6(7):1137–42.

64 Murnane KS. Serotonin 2A receptors are a stress response system: implications for post-traumatic stress disorder. Behav Pharmacol. 2019;30(2 and 3-Spec Issue):151-62.

65 Golden TL, Magsamen S, Sandu CC, Lin S, Roebuck GM, Shi KM, et al. Effects of Setting on Psychedelic Experiences, Therapies, and Outcomes: A Rapid Scoping Review of the Literature. Curr Top Behav Neurosci. 2022;56:35–70.

